# Three dimensional cross-modal image inference: label-free methods for subcellular structure prediction

**DOI:** 10.1101/216606

**Authors:** Chek Ounkomol, Daniel A. Fernandes, Sharmishtaa Seshamani, Mary M. Maleckar, Forrest Collman, Gregory R. Johnson

## Abstract

Fluorescence microscopy has enabled imaging of key subcellular structures in living cells; however, the use of fluorescent dyes and proteins is often expensive, time-consuming, and damaging to cells. Here, we present a tool for the prediction of fluorescently labeled structures in live cells solely from 3D brightfield microscopy images. We show the utility of this approach in predicting several structures of interest from the same static 3D brightfield image, and show that the same tool can prospectively be used to predict the spatiotemporal position of these structures from a bright-field time series. This approach could also be useful in a variety of application areas, such as cross-modal image registration, quantification of live cell imaging, and determination of cell state changes.

## 1. Introduction

A grand challenge of modern cell biology is to understand and model living cells as integrated systems. A critical first step is to determine the localization of key subcellular structures and other functional protein assemblies in living cells. While a variety of microscopy modalities are being used to elucidate the details of cellular organization, they come with trade-offs with respect to expense, spatio-temporal resolution, and cell health. This is particularly acute using fluorescence microscopy, the current method of choice for live cell imaging, wherein samples are subject to phototoxicity and photobleaching, limiting both the quality and time scale of acquisition. Additionally, the number of fluorophores that may be used simultaneously is limited by the number of useful colors, probe perturbation, and spectrum saturation and also requires expensive instrumentation and sample preparation.

In contrast, transmitted light microscopy, e.g., bright field, phase, DIC, etc., is a low-cost alternative modality that greatly reduces phototoxicity and sample preparation complexity. It convolves refractive index differences into a complex image and therefore lacks the readily accessible, organelle identifying advantages of fluorescence microscopy. There exists, however, non-trivial relationships between complimentary modalities that have not yet been exploited. Given spatially registered cross-modal image pairs (i.e. transmitted light and fluorescence microscopy), it may be possible to learn the relationships between the two modalities directly from the images themselves. This knowledge, in turn, can then be applied to new images of the first modality, to predict images of the second. This is particularly attractive when the first image type is inexpensive, less invasive and easy to obtain, as is transmitted light; and the second modality is more costly and complex, e.g., fluorescence microscopy.

Predicting fluorescence microscopy from transmitted light images is a task ideally suited to a subclass of learnable, nonlinear functions known as convolutional neural networks (CNNs). In recent years, they have been used in biomedical imaging for a wide range of tasks, including image classification, object segmentation, and estimation of image transformations. In this study, we first present a tool, based on a widely applied U-Net architecture (Ronneberger et al., 2015), that models cross-modal relationships between different imaging modalities (Figure 1). We demonstrate the utility of this tool for the prediction of 3D fluorescence microscopy images, including detailed labeled structure localization patterns, from 3D transmitted light microscopy images. We do this by quantifying the relationship between transmitted light images and images of the localization of a range of dye and GFP-labeled structures in live cell images. We also demonstrate that the model performs well on 3D time lapse images, and can be used to predict the spatiotemporal position of many subcellular structures from transmitted light images.

**Figure 1.**
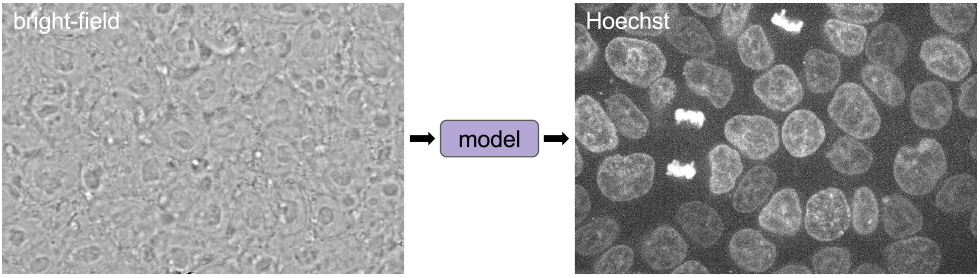
Transmitted light modality images contain information related to alternate imaging modalities. In the above example, we illustrate the need for a to predict a Hoechst staining pattern (for DNA) from a bright-field image.

## 2. Methods

### 2.1. Model description

We employed a network based on various U-Net/V-Net/3D U-Net architectures (Ronneberger et al., 2015; Milletari et al., 2016; Çiçek et al., 2016), first applied towards segmentation tasks in electron microscopy and tissue culture images, due to demonstrated performance in a variety of learning tasks. In practice, it is likely that there exist multiple image-to-image networks that could perform well; a diagram of the model is shown in Figure 2.

**Figure 2.**
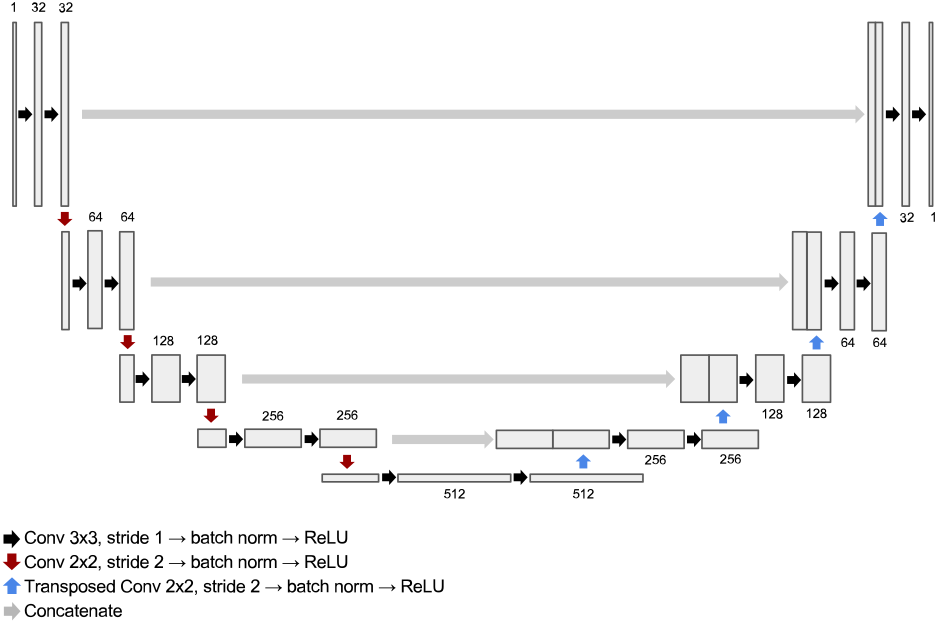
Overview of model. There is no ReLU on the output layer.

Our model uses a fully-convolutional architecture, and is therefore capable of processing input of variable size. This allows us to train a model using multiple small inputs (see 2.3), and perform prediction tasks using full-size images.

The network models a relationship between two microscopy imaging modalities, learned from training on paired, spatially registered images of the same field of cells. Given an input image from one imaging modality, the model outputs a corresponding, predicted, image in the target imaging modality. In this study, we have specifically applied the model to transmitted light microscopy cell images (bright-field and DIC) to predict corresponding fluorescence images, which can then be used e.g. to examine subcellular structures of interest in these cells. However, its use is general and should be readily applicable to other transmitted light modalities, e.g., phase, polarization, reflectance, etc.

### 2.2. Data collection and preprocessing

Our data were derived from a collection of four-channel, 3D z-stacks of human induced pluripotent stem (hiPS) cells from spinning-disk confocal microscopy openly available on the Allen Institute for Cell Science website (allencell.org). The collection consists of high replicate 3D z-stacks of genome-edited hiPS cell lines, in each of which a particular subcellular structure was endogenously tagged with a molecule fused to GFP (Roberts et al., 2017). The 4 channels include: (1) a transmitted light modality image (either bright-field or DIC) and fluorescence images of either (2) a fluorescent dye that labels the cell membrane (Cell-Mask), (3) a fluorescent dye that labels DNA (Hoechst), and (4) the endogeously mEGFP-tagged subcellular structure.

For static z-stacks, we created paired-image data sets by pairing the transmitted light image (channel 1) with a corresponding fluorescence channel (channels 3 or 4). These were then used to train the CNN to learn the relationship between the transmitted light images and fluorescence images of a particular subcellular structure.

For time series experiments, we generated new time lapse z-stacks of the same field of cells taken every 35 seconds at a magnification of 0.065*μ*/px (lateral scale), and used the model to predict the location of the structure. A full summary of the data sets used to train and test the models are in Table 1.

**Table 1.** Summary of image pairs used in this manuscript.

| Input | Fluorophore | Labeled structure |
| --- | --- | --- |
| bright-field | Hoechst | DNA |
| bright-field | CellMask | plasma membrane |
| DIC | CellMask | plasma membrane |
| bright-field | mEGFP α-tubulin | microtubules |
| bright-field | mEGFP β-actin | actin filaments |
| bright-field | mEGFP Desmoplakin | desmosomes |
| bright-field | mEGFP Fibrillarin | nucleoli |
| bright-field | mEGFP Lamin B1 | nuclear membrane |
| DIC | mEGFP Lamin B1 | nuclear membrane |
| bright-field | mEGFP Myosin IIB | Actomyosin bundles |
| bright-field | mEGFP Sec61β | endoplasmic reticulum |
| bright-field | mEGFP TOM20 | mitochondria |
| bright-field | mEGFP ZO1 | tight junctions |

All input and target images (see 2.3) were pre-processed by z-scoring pixel intensities across each image. We do not perform any data augmentation procedures on our training data, including padding, flipping, or warping.

### 2.3. Training procedure and implementation

We trained a model for each collection of input-target image pairs (Table 1). This resulted in 13 independent models. We used 30 image-pairs for the training set and allocated all of the remaining image pairs to the test set. The images were resized such that each voxel corresponds to a 0.3x0.3x0.3um cube. Due to current memory constraints associated with 3D convolution and our current GPU computing configuration, we were not able to train the model directly on the full-sized images.

Considering these practical constraints, we trained the model on batches of 3D patches, subsampled uniformly (both across all training images as well as spatially within an image) from training images at random. The training procedure takes place in a feed-forward fashion typical of these methods, as described in Algorithm 1. See our software repository (4) for further details.

We can interpret the models’ generalization error as an estimate of the relationship between input and target images. We report this error as the L2 loss between a predicted image and its target image, normalized such that the prediction of a blank (all pixel intensities zero) image would yield an error of 1.0.

Our model training pipeline was implemented in PyTorch (http://pytorch.org). Each model was trained using the Adam optimizer (Kingma & Ba, 2014) with a learning rate of 0.001 for 50,000 mini-batch iterations of size 24. Running on a Pascal Titan X, each model completed training in approximately 16 hours.

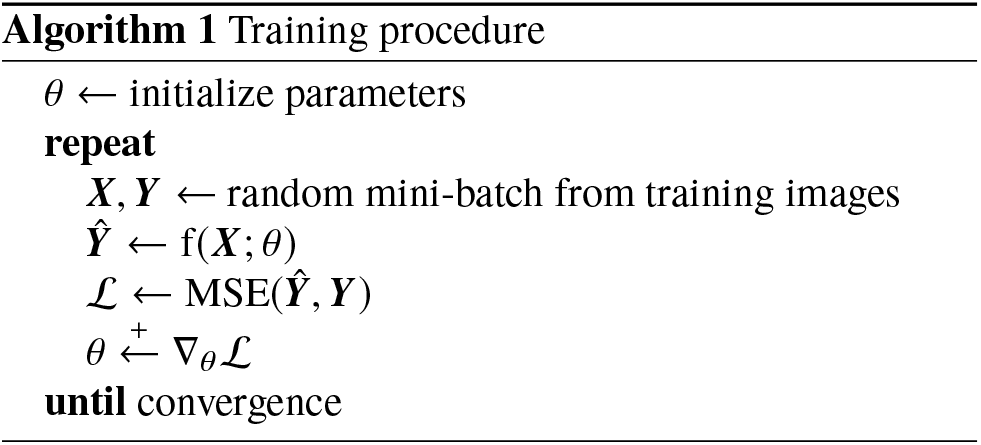

## 3. Results

### 3.1. Relationship between input images and predicted fluorescence images

Figure 3 demonstrates the application of our model to a 3D bright-field image to predict a 3D fluorescence image of Hoechst dye localizing to DNA. The predicted images demonstrate good qualitative correspondence with target fluorescence images in all z-slices. The individual nuclear regions are well-formed and separated in the predicted image, and it is possible to distinguish between the DNA of mitotic and non-mitotic cells. Lamin B1, a nuclear marker, also tags the nuclear region as well and reorganizes during mitosis (Figure S2). Other example predictions include bright-field to fibrillarin, bright-field to Tom20, and DIC to LaminB1 (Figure 4a). Predictions for all the labels used in this manuscript are shown in supplementary figures S1 to S12.

**Figure 3.**
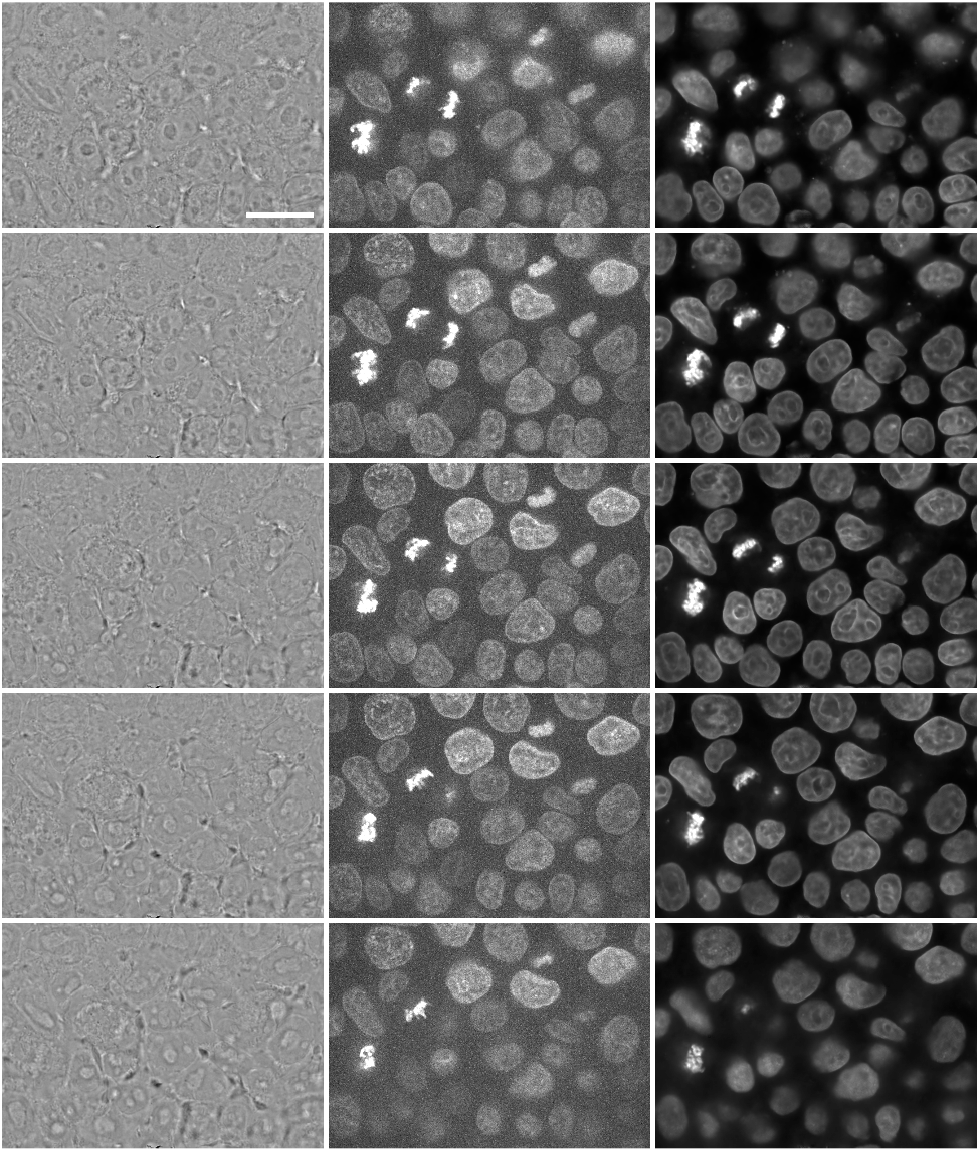
Predicted Hoechst fluorescence image from bright-field input image. Left column shows z-slices of a 3D bright-field image at 1.5*μ*m intervals moving top to bottom of the z-stack (input image). Middle column shows z-slices from the corresponding observed Hoechst fluorescence image (target image). Right column shows the model-predicted Hoechst fluorescence image (output image). Scale bar is 20 *μ*m.

**Figure 4.**
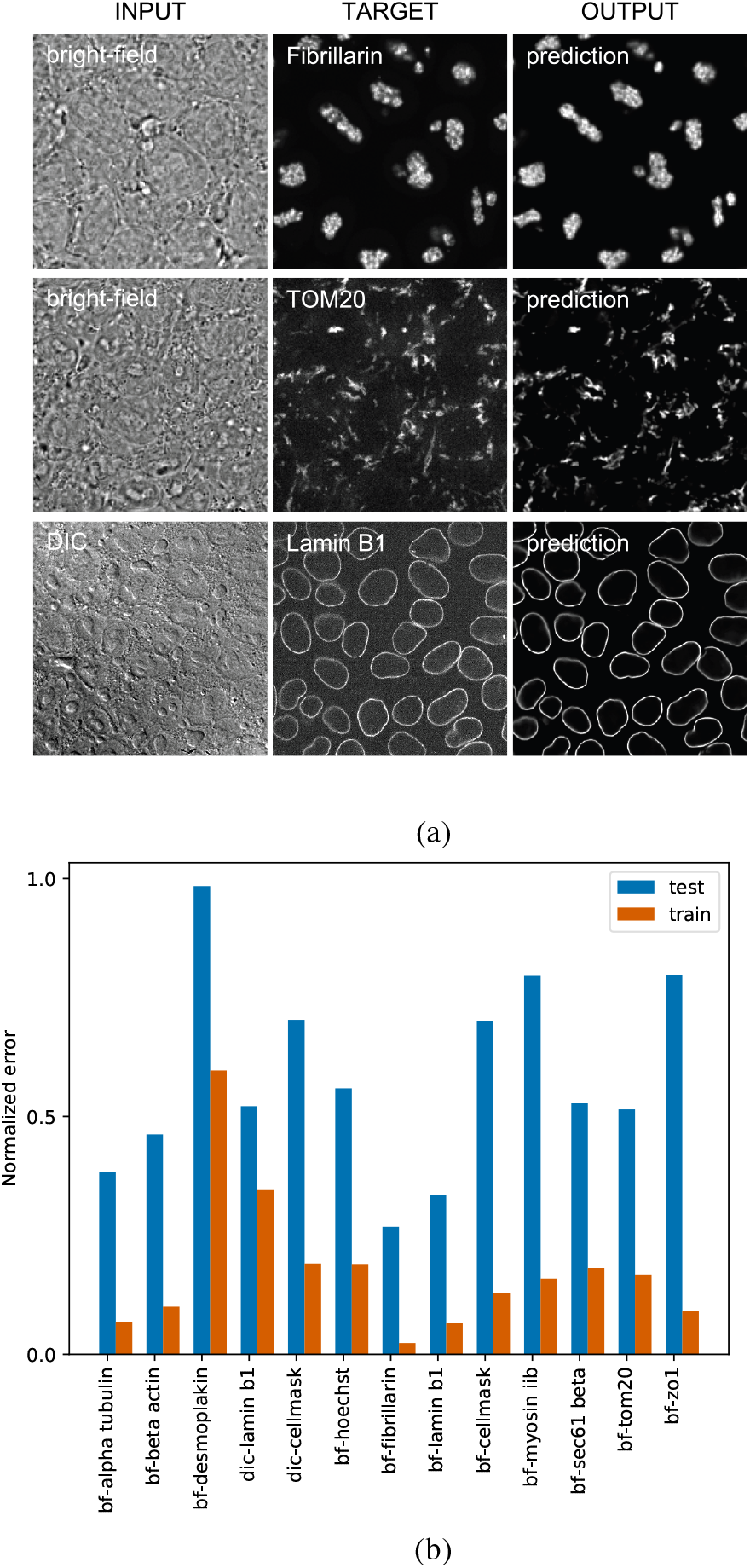
(a) Example input images (left column; showing middle slice), target images (middle column) and predicted images (right column). (b) Train and test set errors for the complete image collection.

We have assessed performance of the models by a normalized training loss function as described above (2.3); corresponding errors for training and test sets for all 13 models are shown in Figure 4b) This estimated performance depends on the target subcellular structure; for example, nuclear structures (fibrillarin, Lamin B1, and Hoechst) perform comparatively well, while e.g. the desmoplakin model performs relatively poorly, likely due to a lack of contrasting features in the bright field image and perhaps the poor signal to noise (Figure S1); tubulin (microtubules, S3) also did not perform well in the model probably also because of the absence appropriate features in the bright field image.

### 3.2. Combined, multiple-structure predictions

We tested the feasibility of using conjoining models to generate multi-structure predicted images. In Figure 5a, we applied an identical input image to each of the trained structure prediction models in the collection and combined the outputs into a single, multi-structure predicted image. In Figure 5b, we show the multi-structure result: predicted fluorescence images from top-performing models given the same input bright-field image (bright-field to Hoechst, Lamin B1, Sec61, fibrillarin, and Tom20). The model outputs can then be combined into a single, multi-structure visualization, as well as potentially combined with actual fluorescence data, to augment the effective number of available channels (Figure 5b) in an imaging experiment.

**Figure 5.**
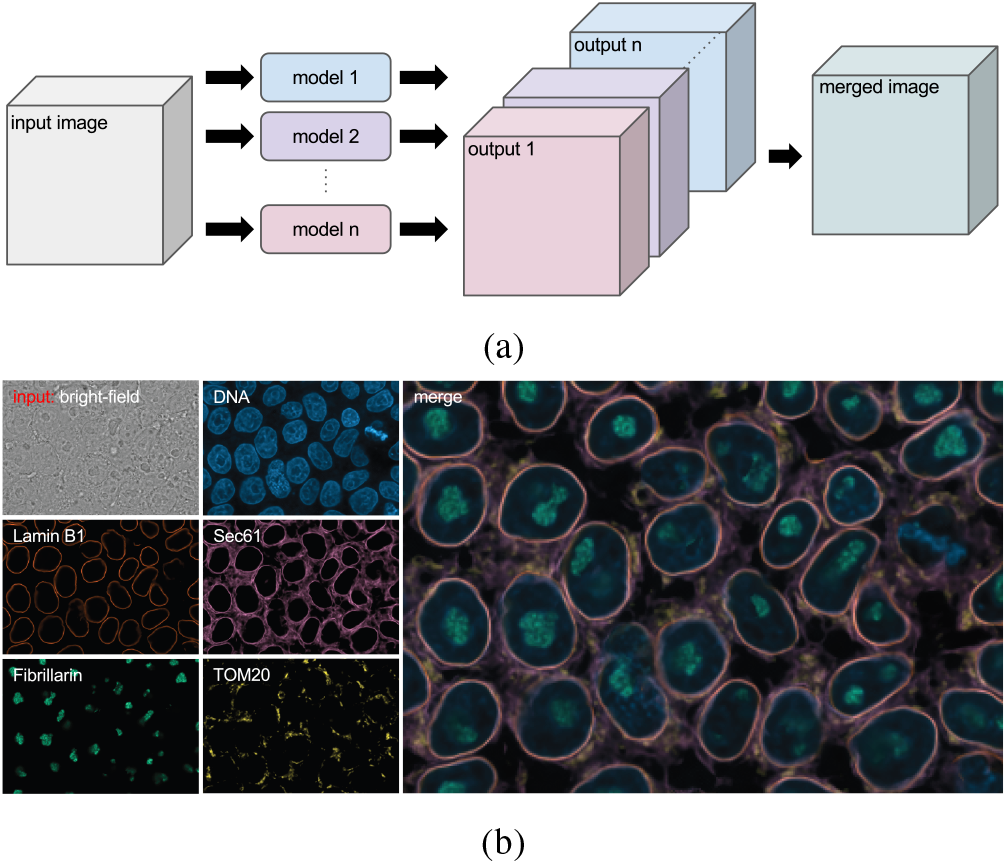
(a) Multiple structure prediction models are applied to the same input image, resulting in an output image per model. Those are combined to form a multi-color image. (b) Example input image (showing middle slice) and corresponding predicted structures are merged.

### 3.3. Time series

We also asked whether the trained model collection could be used to predict the dynamics of a temporal fluorescence image sequence. We did this by first applying each frame of a 3D bright-field time series to the model collection and then combining these outputs into video. The results suggest that models of this type, trained only on a sequence of static image pairs, can be used to predicting time-series for which no fluorescence imaging target is available. Center slices of serial time points are shown in Figure 6. The corresponding video is available at https://youtu.be/wObyJASI574.

**Figure 6.**
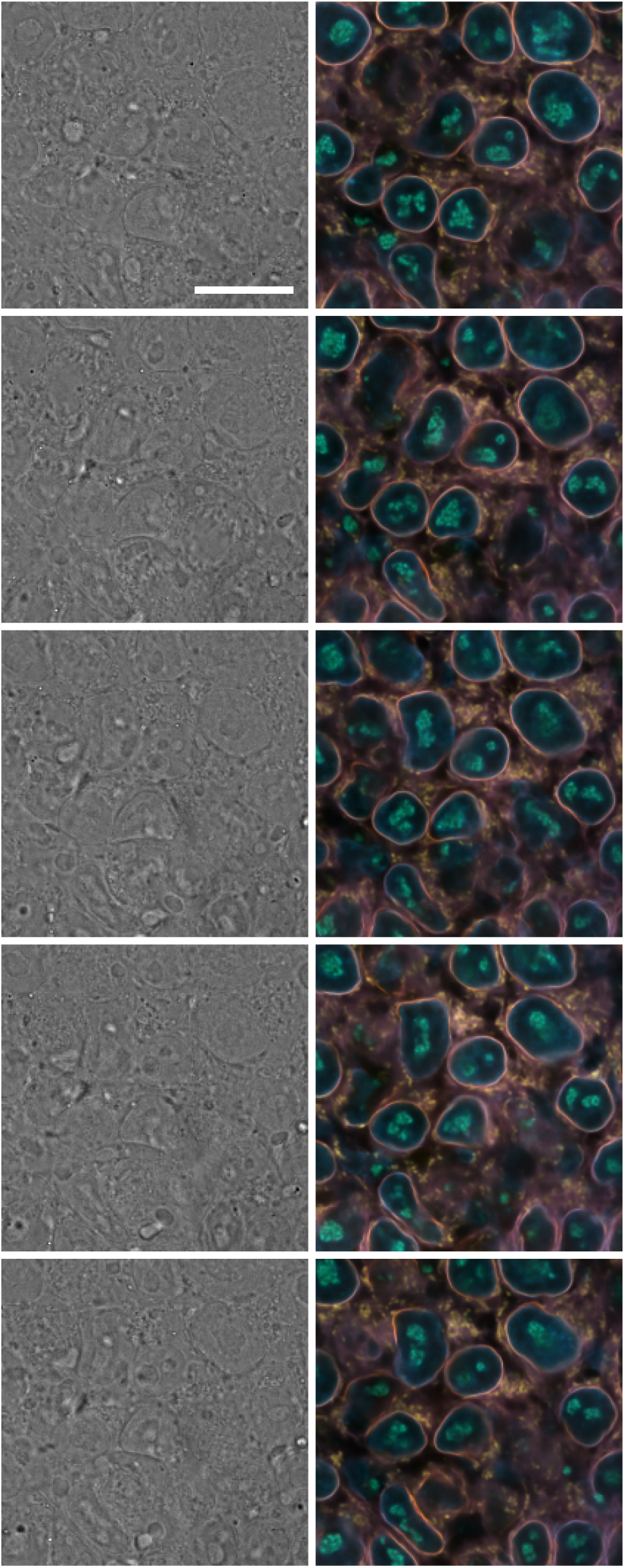
Combined structure predictions on time-series image set. Each row is a separate time point with an interval of 280 seconds between time points. Time advances from top to bottom. Left column: single z-slices from the input bright-field z-stacks. Right column: merged corresponding predicted fluorescence images from Hoechst, Fibrillarin, Lamin B1, Sec61, and TOM20 models. Scale bar is 20 *μ*m.

This proof-of-concept demonstrates potential model utility for prediction on input images acquired in a variable fashion to those images on which the models are trained. Specifically, in this case, the time series images were acquired at a different magnification: 0.065 /px for the time-series versus 0.108 /px for all static training image data sets, and therefore possess a different lateral resolution. Nevertheless, the predicted time series images were of sufficient quality to visualize multiple cellular structures from live cells over a long period of time, implying potential model robustness.

## 4. Discussion

We have introduced a method for cross-modality microscopy image prediction based on an 3D convolutional neural network architecture. We have shown how, once trained on paired image sets, this tool can successfully predict 3D fluorescence images from transmitted light microscopy images. While much remains to be done to extend and further validate the tool, these initial observations are very promising. Of course, biological samples in a typical research imaging setting may have significant sample and image variability. However, the model appears robust as seen in its performance with time-series data, which predicted time lapse fluorescence from input data of different resolution, modality, and timing than that on which the model was trained. In summary, the presented tool offers the potential for biologists to take advantage of fluorescence techniques via prediction, while chiefly producing only resource-friendly transmitted light images. Moreover, direct modeling - mapping one image modality to another - paves the way for evaluation of the organization of several subcellular structures, simultaneously, effectively offering multi-channel fluorescence results from a single transmitted light image.

Despite the potential advances introduced by this tool, the method still presents several limitations. Even given training images that are of high quality and a model of sufficient power, the current method will perform poorly if the features in the transmitted light image do not provide sufficient information for a correlative, learnable relationship between the input and target images. In this context, the model does well with membranous structures (ER, mitochondria, nucleus) and larger aggregates (DNA, nucleoli), it does not appear to do as well as structures like desmosomes or microtubules. Additionally, the model may be limited in its ability to predict relationships for which there are few or no examples in the training data. For example, early experiments with our model failed to accurately predict DNA localization during mitosis, as initial training images included few mitotic cells.

Model performance may be further optimized by applying a variety of model architectures, and/or tuning model hyperparameters, as is common in neural network development, though we have not focused on these detailed engineering aspects here. There are also multiple potential sources of model error. The most apparent is the quality of the imaging used to train and apply the model. Camera noise is one example; for example, in the case of Hoechst prediction (see Figure 3), the output image seems to be an “average”, de-noised version of the target image. It should be noted that we were able to achieve good generalization performance with the handful of training images used. The quantity of data used here is much less than is typical in deep network training applications, and our generalization performance is likely to improve with more data.

While our methodology can be used to directly evaluate relationships among imaging modalities, to predict images of one modality from another, and to subsequently visualize localization patterns as described above, it may also be suitable for a variety of other tasks. One straightforward application is to integrate model fluorescent label predictions into pre-existing fluorophore-dependent image processing pipelines to perform tasks such as dense 3D cell segmentation or cross-modality image registration. With model-predicted images of sufficient quality, it may be possible to reduce or even eliminate the need to routinely capture fluorescence images in imaging pipelines, permitting the same throughput in a far more cost-effective and efficient manner. Furthermore, other transmitted light modalities, like polarization, may reveal new, learnable features. Finally, when trained with fluorescent proteins that reveal details within an organelle like the nucleus, for example, it may be possible to see changes in activities that accompany that accompany cell state transitions.

### Software and Data

Sofware for training and using models is available at https://github.com/AllenCellModeling/pytorch_fnet. The data used to train the model is available at http://www.allencell.org.

## Acknowledgements

We thank the members of the Allen Institute for Cell Science team, who generated and characterized the gene-edited hiPS cell lines, developed image-based assays, and recorded the high replicate data sets suitable for modeling. We would like to especially thank the AICS Microscopy group for providing images of different transmitted-light imaging modalities. These contributions were absolutely critical for model development.

We thank Eric Christiansen and Phillip Nelson for discussions on 2D deep learning.

We thank Paul G. Allen, founder of the Allen Institute for Cell Science, for his vision, encouragement and support.

## Author Contributions

GRJ conceived the project. DAF and GRJ demonstrated proof of concept for 2D images. CO implemented the model for 3D images. CO, DAF, SS, FC, and GRJ designed experiments and wrote the paper, MM provided guidance and helped write the paper.

---

**Figure S1.**
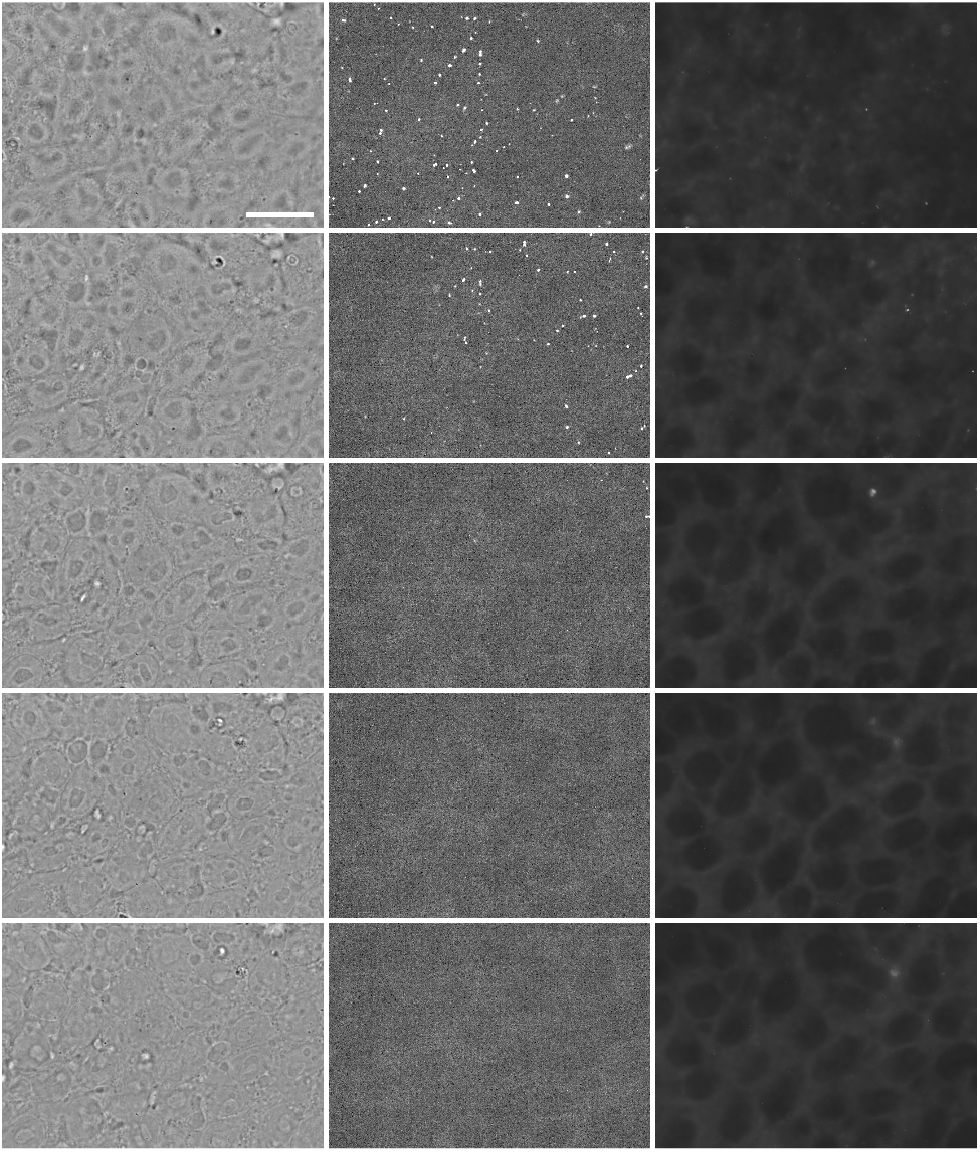
Predicted Desmoplakin fluorescence image from bright-field input image. Left column: z-slices of an input 3D image at 1.5 *μ*m intervals from top to bottom of the z-stack (input image). Middle column: z-slices from the corresponding observed target fluorescence image. Right column: z-slices from the model-predicted fluorescence image. Scale bar is 20 *μ*m.

**Figure S2.**
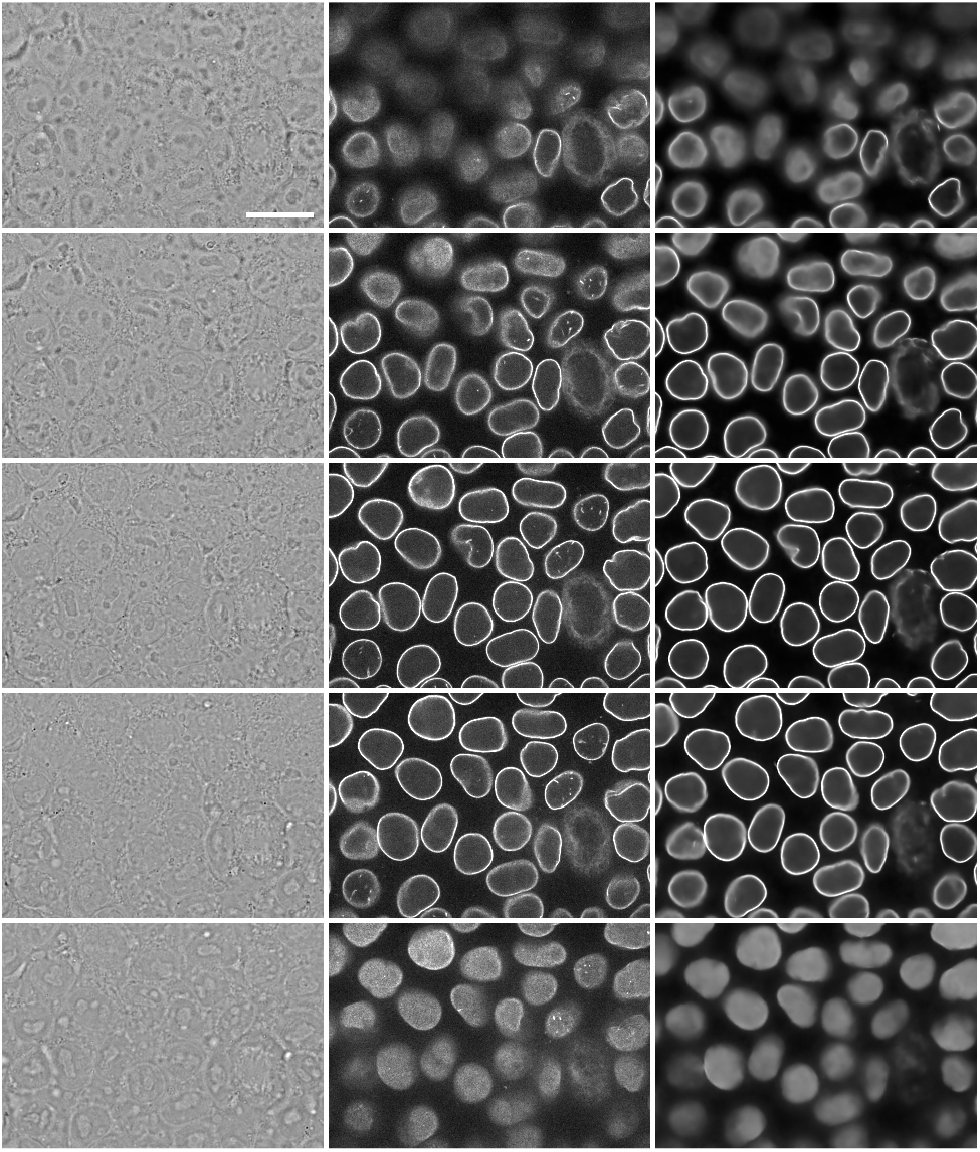
Predicted Lamin B1 fluorescence image from bright-field input image. See figure S1 caption for column descriptions.

**Figure S3.**
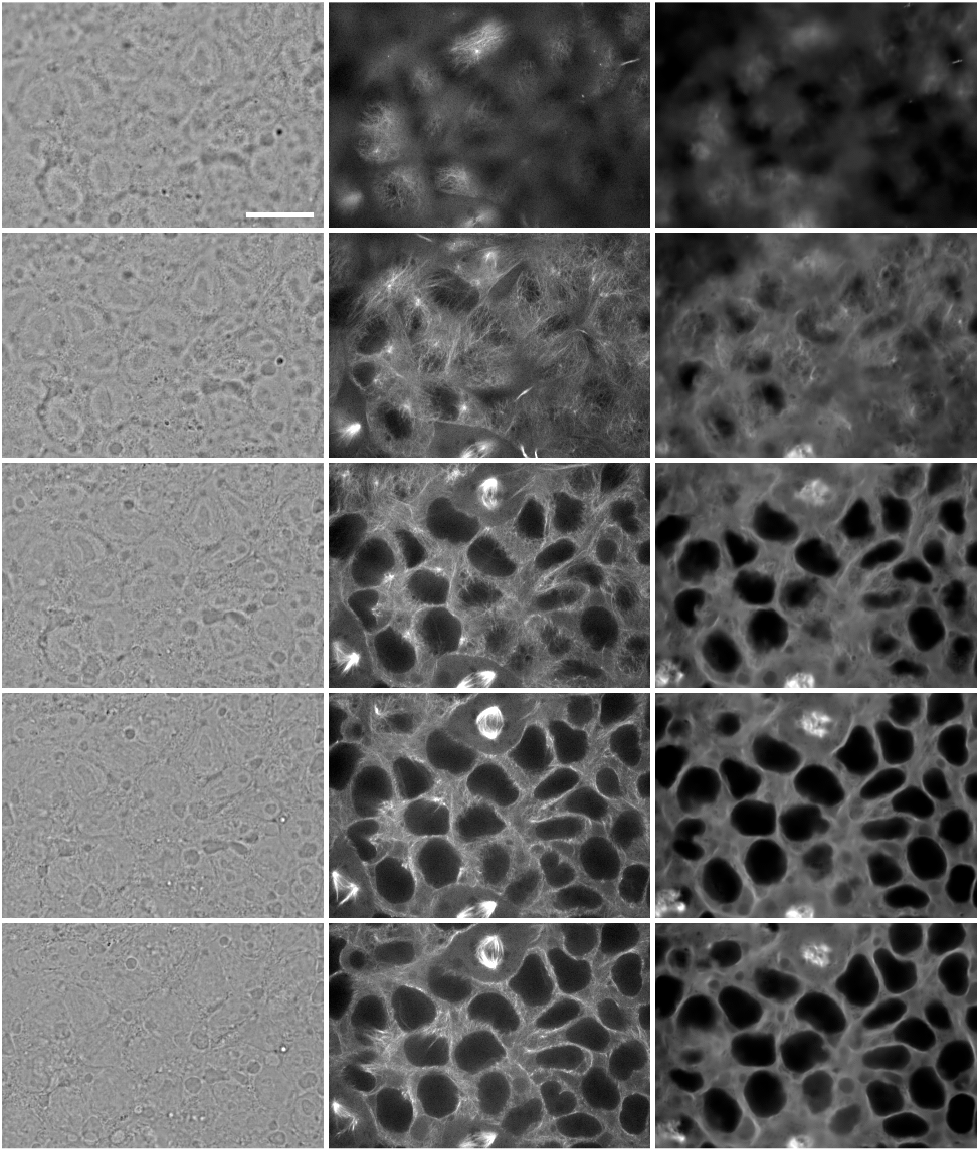
Predicted α-tubulin fluorescence image from bright-field input image. See figure S1 caption for column descriptions.

**Figure S4.**
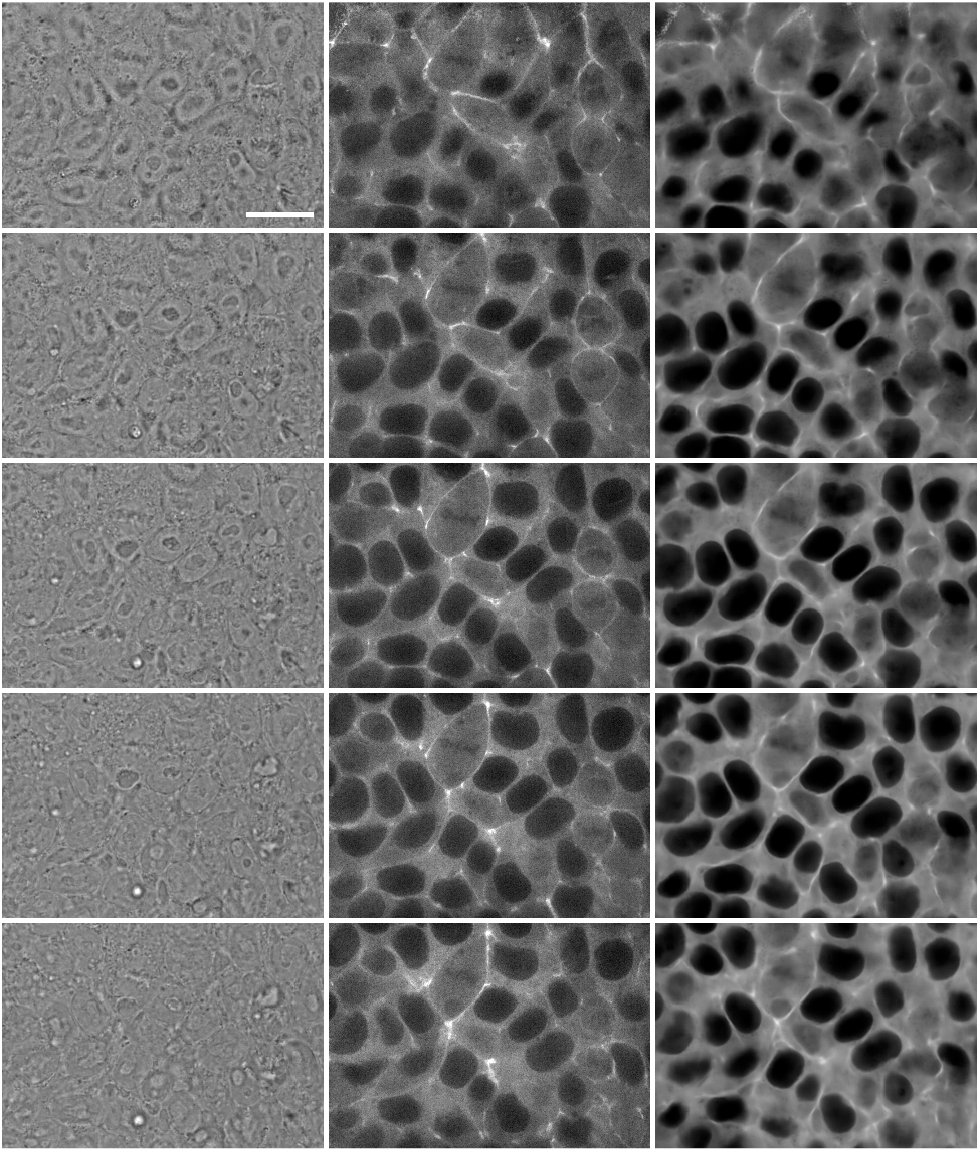
Predicted β-actin fluorescence image from bright-field input image. See figure S1 caption for column descriptions.

**Figure S5.**
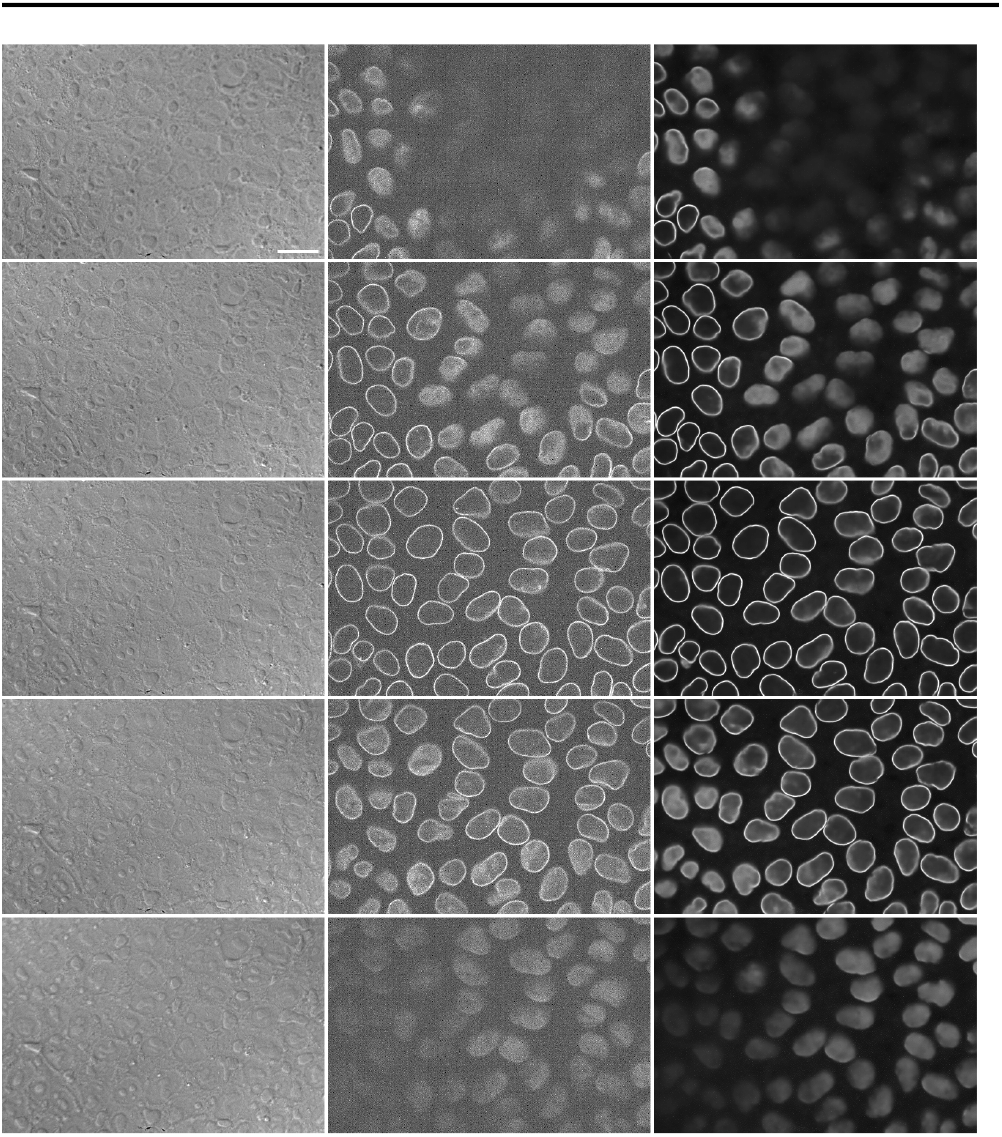
Predicted Lamin B1 fluorescence image from DIC input image. See figure S1 caption for column descriptions.

**Figure S6.**
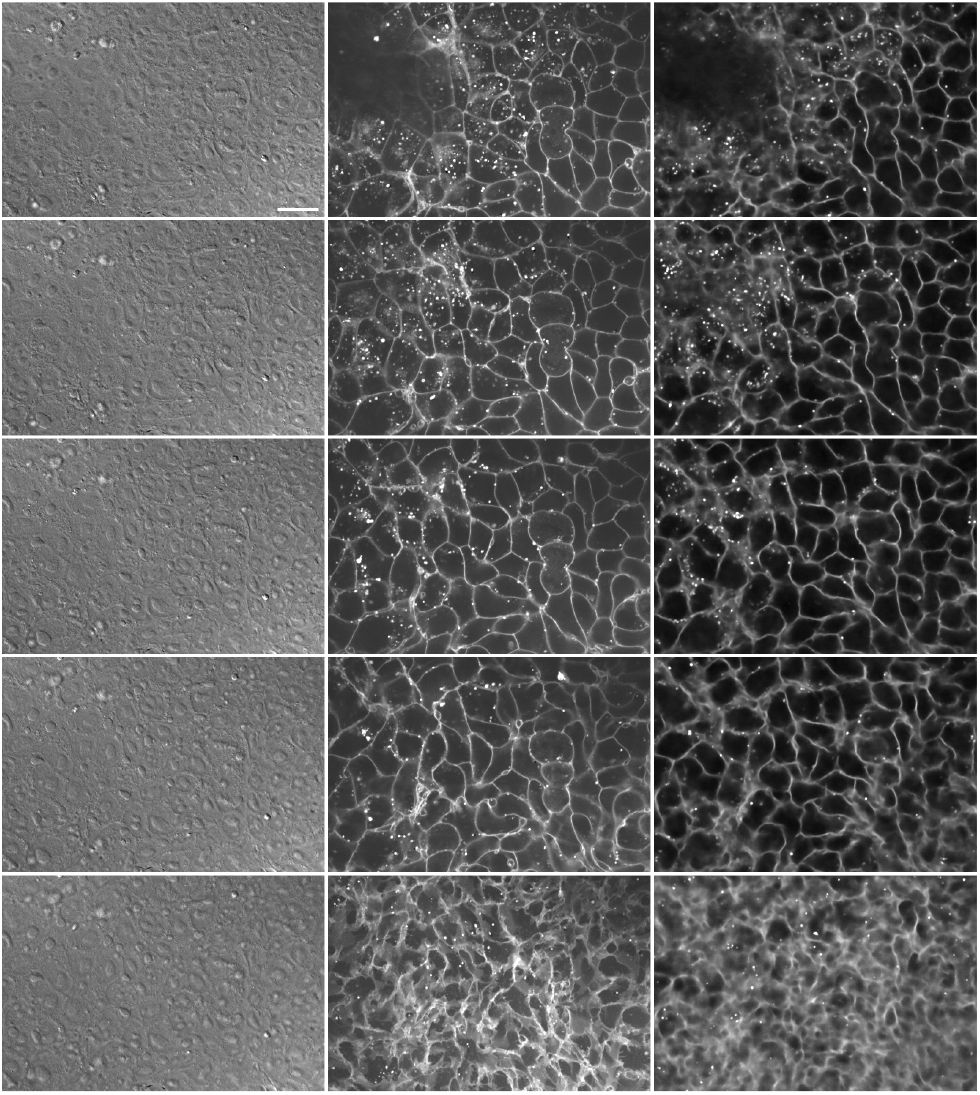
Predicted CellMask fluorescence image from DIC input image. See figure S1 caption for column descriptions.

**Figure S7.**
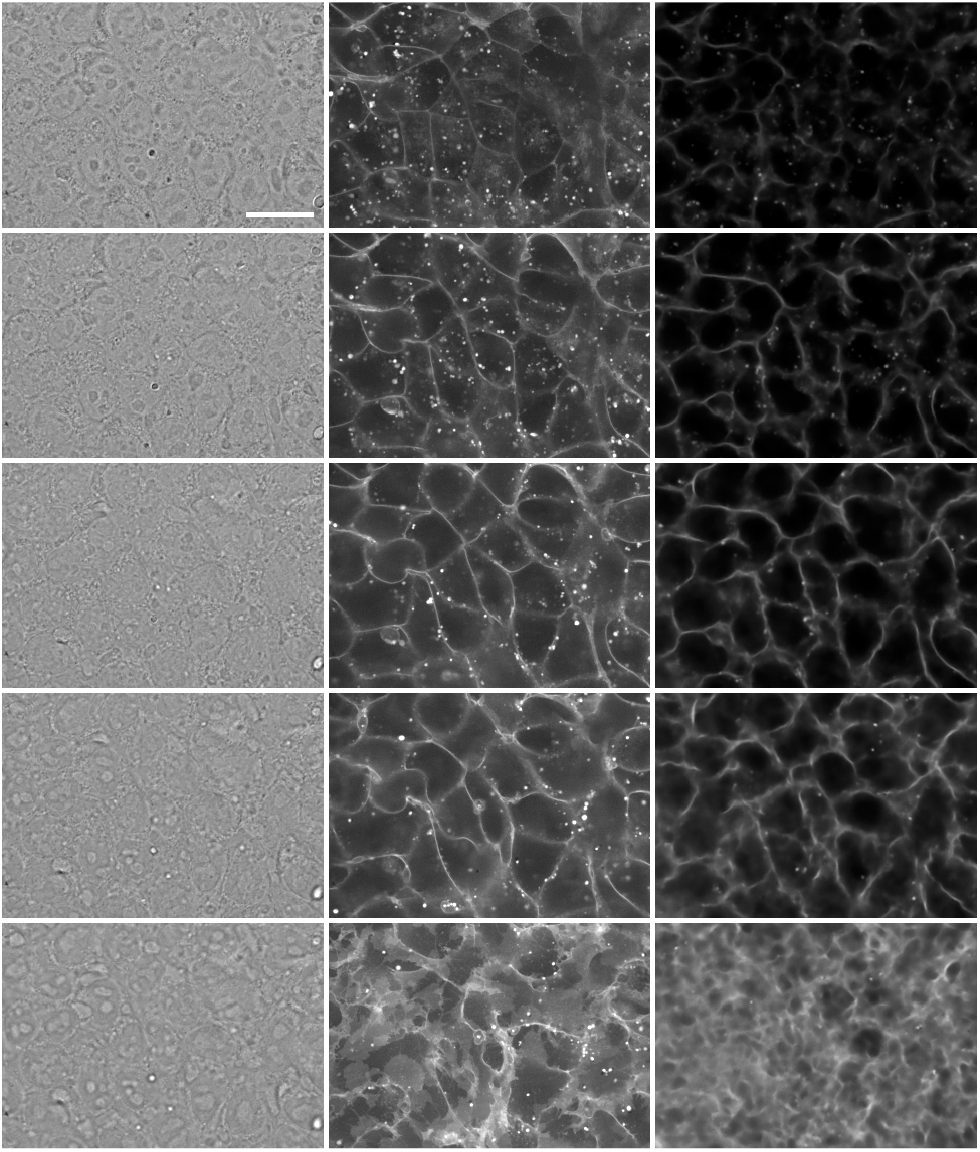
Predicted CellMask fluorescence image from bright-field input image. See figure S1 caption for column descriptions.

**Figure S8.**
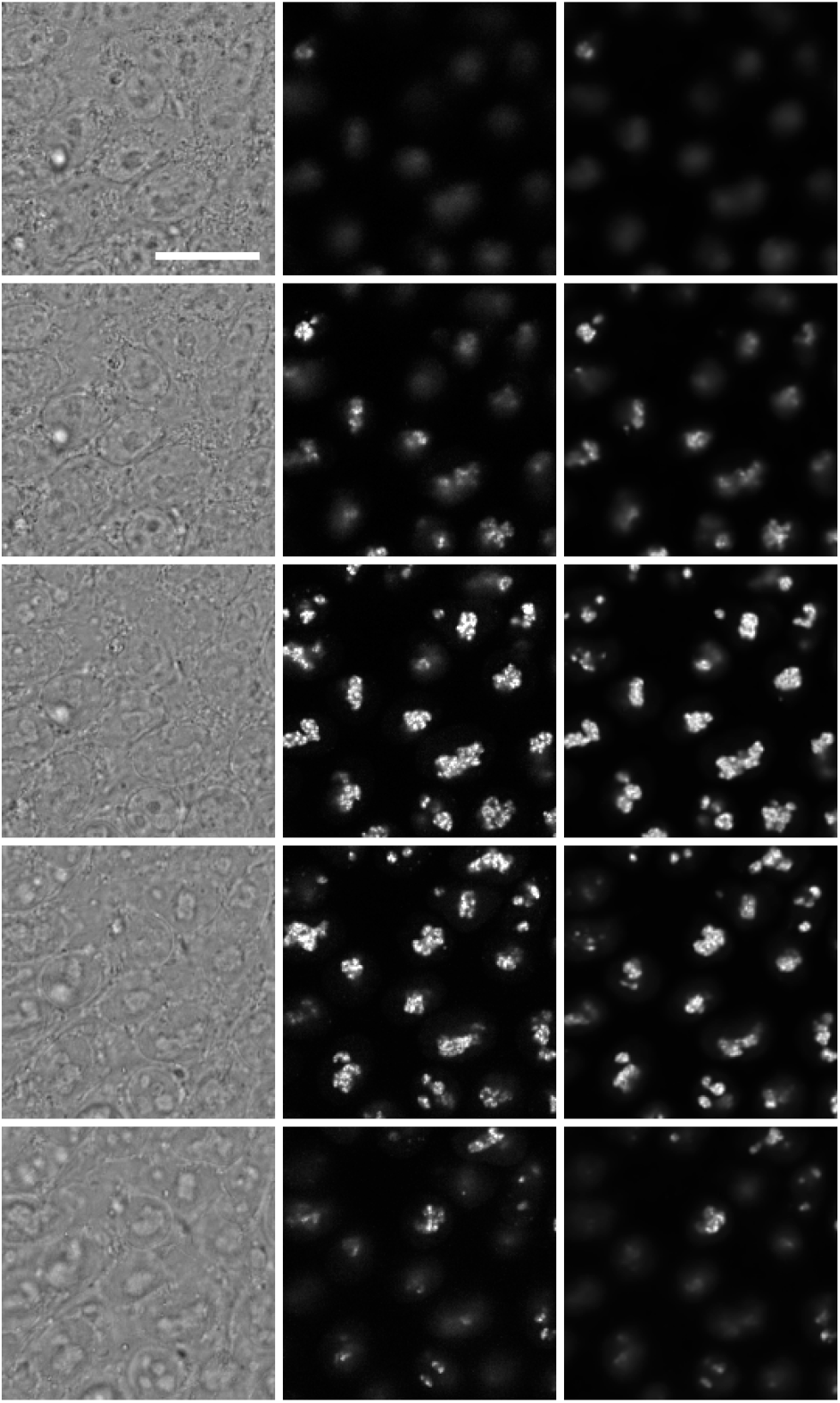
Predicted Fibrillarin fluorescence image from bright-field input image. See figure S1 caption for column descriptions.

**Figure S9.**
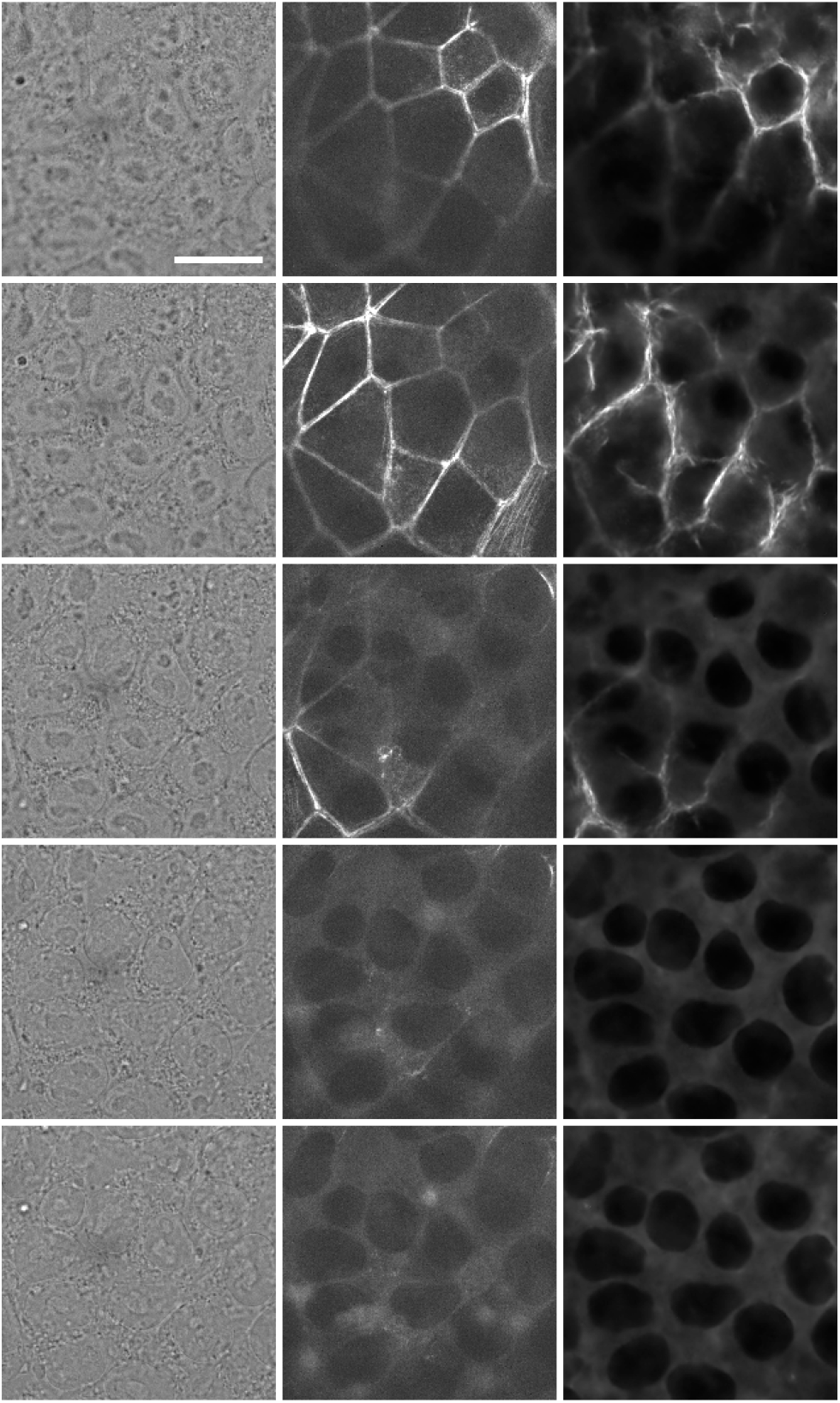
Predicted Myosin IIB fluorescence image from bright-field input image. See figure S1 caption for column descriptions.

**Figure S10.**
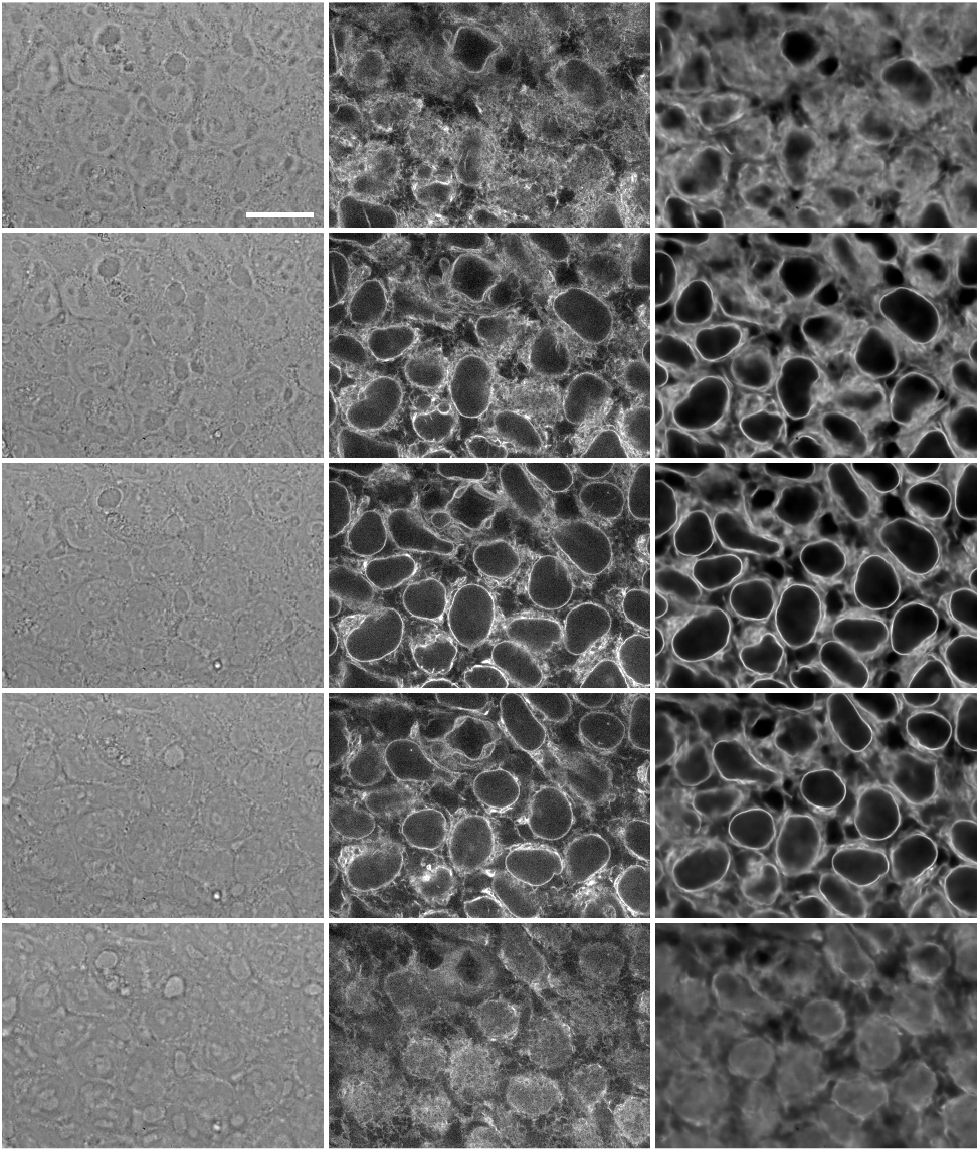
Predicted Sec61 βfluorescence image from bright-field input image. See figure S1 caption for column descriptions.

**Figure S11.**
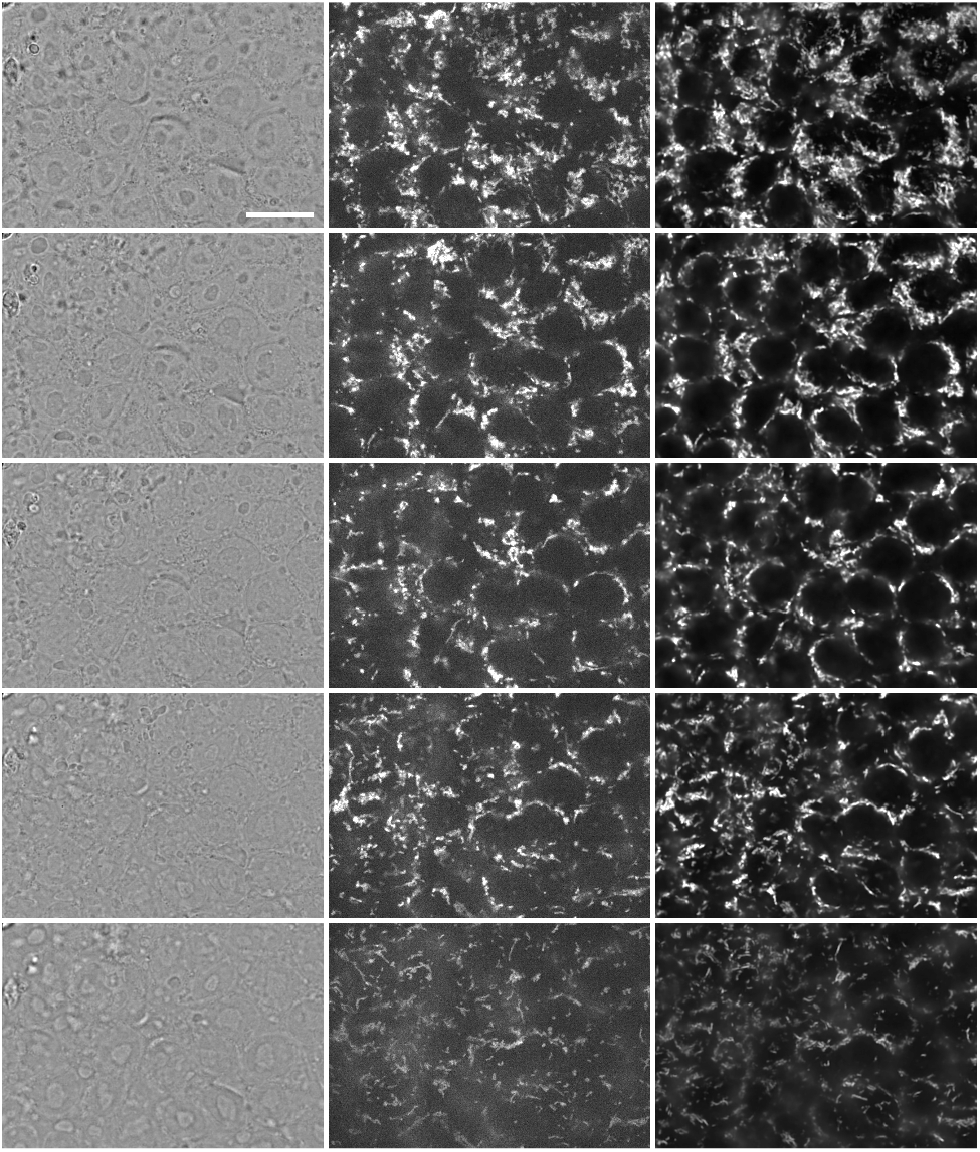
Predicted Tom20 fluorescence image from bright-field input image. See figure S1 caption for column descriptions.

**Figure S12.**
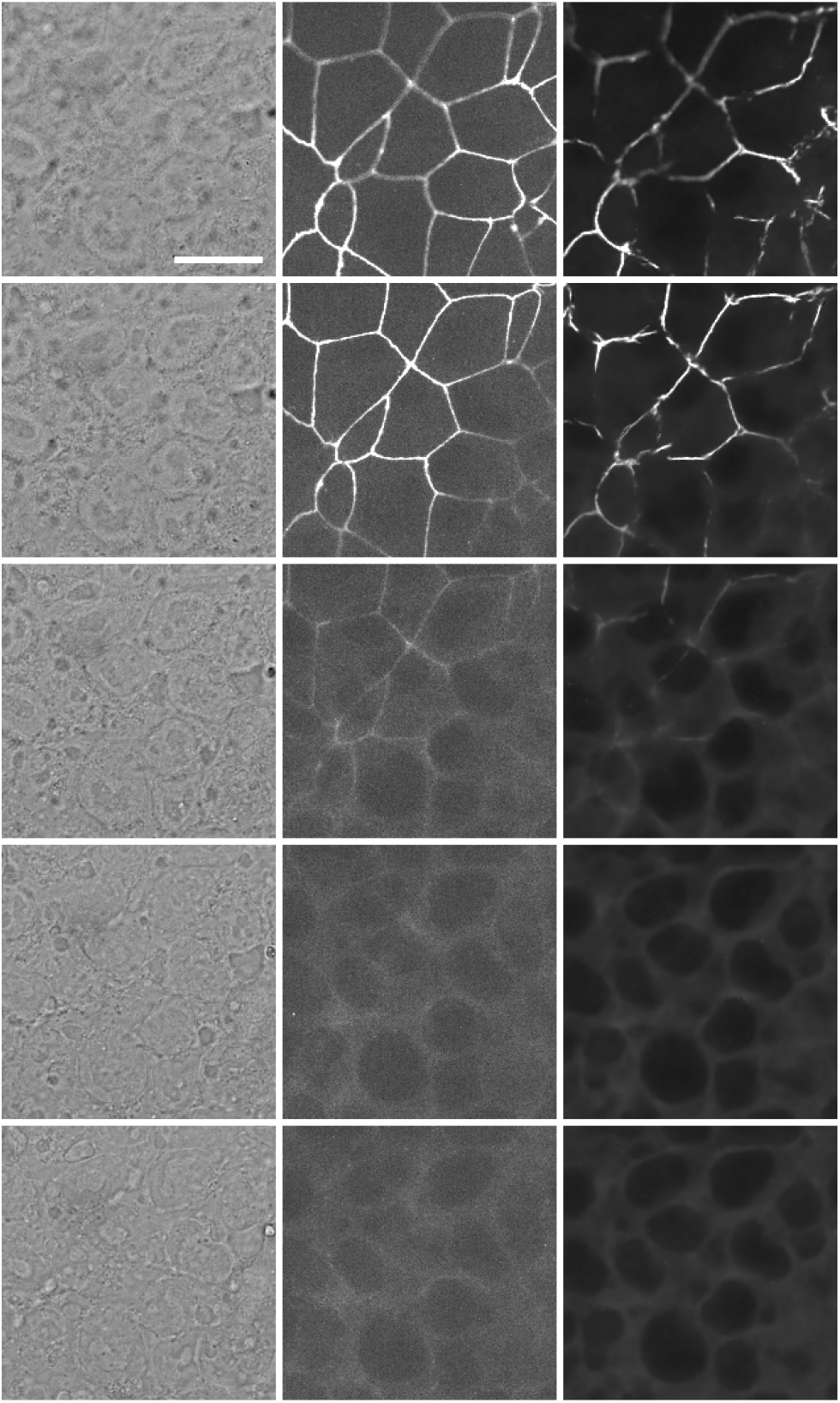
Predicted ZO1 fluorescence image from bright-field input image. See figure S1 caption for column descriptions.

